# Optogenetic construction of *de novo* integrin-adhesion complexes reveals role for biocondensation in adhesion nucleation

**DOI:** 10.1101/2025.10.08.681107

**Authors:** Minggang Xiong, Tsun Lam Leong, Mengyu Chen, Jiabei Chen, Tsun Chiu Lee, Cheng-Han Yu, Artem K. Efremov, Heath E. Johnson

## Abstract

Integrin-adhesion complexes (IACs) form spontaneously in cells on extracellular matrix substrates, allowing them to sense matrix composition and transduce force. However, IACs often do not form uniformly across a cell, which begs the question: What is required to nucleate an adhesion, and what factors enable the stabilization of an IAC once it has formed? Many factors have been suggested to promote formation and the subsequent stabilization of IACs. It is difficult to explicitly test these factors *in vivo* as IACs undergo constant remodeling. Here, we employ optogenetics to explicitly test the ability of talin in different activity and phase states to nucleate and stabilize IACs in regions where none are present. We find that fusion of intrinsically disordered regions directly to talin enhances its adhesion nucleation potential and allows new adhesions to be produced in response to optogenetic talin clustering. Similarly, expression of factors previously shown to enhance biocondensation *in vitro*, such as paxillin, the paxillin N-terminus, or unfolding of talin, allows for adhesion nucleation and biocondensation of talin. We show that these biocondensates of talin can cluster and activate integrins even in the absence of extracellular matrix. By applying optogenetic activation to regions of the cell with or without ventral actomyosin, we demonstrate actomyosin engagement promotes the formation and stability of adhesions. These results are corroborated by theoretical modelling which shows that phase separation of talin is enhanced by differential clutch formation in the presence of actomyosin thus enabling peripheral adhesion formation and stability. This work establishes a model in which increased cooperativity of talin enables IAC nucleation through talin biocondensation, which clusters and activates integrins. In addition to these findings, we generate multiple optogenetic tools that enable local nucleation or enhancement of IACs.

**Highlights:**

1. Optogenetic tools mediating talin biocondensation can locally induce focal adhesion formation
2. Paxillin LD domains enable biocondensation of talin
3. Biocondensation of talin enables IAC formation
4. Phase separation of talin can activate integrins independently of ECM
5. Computational model reconciles spatial variance in LLPS and IAC formation.

## Introduction

The linkage between cells and the underlying extracellular matrix (ECM) is an essential component of many tissues and a mediator of morphogenesis and cell migration^1,2^. This role is filled by integrin-adhesion complexes (IACs), which form a physical linkage between the ECM and the actin cytoskeleton.^3^ At the heart of IACs are clusters of integrins, heterodimeric membrane receptors that bind specific ECM proteins outside the cell and link to adaptors on their cytosolic side.^4^ A key adaptor protein responsible for bridging the gap between integrins and the cytoskeleton is talin. Talin consists of an integrin binding FERM domain head and a large, flexible rod domain tail and is an essential mediator of integrin activity and adhesion formation. It binds to integrins on their cytoplasmic tails, both activating integrins^5^ and providing a scaffold for many IAC components that will link the integrins to the actin cytoskeleton. In the cytosol, talin is typically in a folded, autoinhibited conformation, maintained by an interaction between the head and rod.^6^ Inhibition can be released through unfolding mediated by phospholipid binding^7^ to the head or possibly through interactions with other adhesion proteins such as vinculin.^8^ As the IAC is exposed to increasing force, talin unfolds, exposing new binding sites and recruiting additional proteins^9^. Thus, talin is a mechanosensitive organizing center in the formation of adhesion complexes.

IACs form spontaneously on cells plated on ECM and are often around the periphery of the cell, where ventral actin is readily available and frequently making new contact with the matrix as the cell protrudes forward. Traction forces are typically highest^10^ around the edge of the cell, which stabilizes the active conformation of the integrins^11^. While retrograde flow may stabilize more mature adhesions linked with actin, nascent adhesion disassembly is accelerated by force.^12^ The number and distribution of IACs vary significantly between cells and contexts, and thus, why IACs form in some places but not others, as well as what it takes to nucleate a new adhesion, remains unclear. The classic model of adhesion nucleation begins with integrin activation, either by talin recruitment or by spontaneous matrix binding. Immobilized integrins then cluster, recruiting various adhesion proteins, linking with actin, and transducing force.^3^ However, it is unclear how and when a single integrin activation event is transitioned to a large adhesion complex made of hundreds of proteins. Even nascent adhesion nanoclusters contain about 50 integrins, as well as many copies of several adaptors such as talin and paxillin.^12^

One possible explanation for how integrins and adhesion adaptors complex together to nucleate adhesions is liquid-liquid phase separation (LLPS), which has emerged in the past decade as playing a central role in the subcellular organization of cells. Weak, multivalent interactions, often through the intrinsically disordered regions (IDRs) of proteins, enable LLPS of proteins and RNA into what is known as biomolecular condensates.^13^ Biomolecular condensates are particularly useful for cellular organization as allow for localized high concentrations of molecules, while still allowing them to be dynamic. Recent work has shown LLPS of various proteins involved in IAC formation occur *in vitro*, including integrins^14^, talin^15^, p130Cas^16^, focal adhesion kinase (FAK)^16^, paxillin^14,17^, LIMD^18^, KANK^19^, tensin^20^, and kindlin^16^. Furthermore, adhesions display many of the properties of LLPS *in vivo*, such as fusion, responses to changes in pH or temperature, and rapid turnover.^21–23^ The precise role of phase separation in *in vivo* IACs is still being defined and has been suggested to enable force sensing^18^, promote integrin clustering^15,16^, IAC formation^16,17^, regulate protein mobility^24^, and even regulate mRNA translation^25^.

Here we sought to understand the requirements to nucleate IACs and to determine whether LLPS can enable IAC formation. To accomplish this, we developed optogenetic tools that would allow us to spatiotemporally control the clustering and localization of talin in the presence or absence of factors which might promote biocondensation. We utilized the *Arabidopsis* photoreceptor cryptochrome 2 (C2), which undergoes tetramerization under blue light^26,27^. In addition to homotetramerizing in response to light activation, C2 can simultaneously form a heterodimer with the N-terminus of *Arabidopsis* CIB1 (CIBN).^28^ By tethering CIBN to the plasma membrane, C2 can be simultaneously recruited to the membrane and tetramerized. Finally, by attaching IDRs to C2, phase separation can be induced on the membrane in a light-controlled manner.^29^ We employed this strategy by fusing different forms of talin to these optogenetic domains to enable us to locally control the degree of cooperativity between talins in response to light.

Utilizing these optogenetic tools, we probe the ability of talin in various states to form new IACs. Our results show that biocondensation of talin - either through paxillin or through unfolding of talin itself - enables integrin clustering, activation, and thus IAC nucleation. LLPS of talin localizes integrins to clusters, while talin recruitment in the absence of sufficient phase separation promoting domains can participate in and enhance existing adhesions but cannot nucleate new ones. Furthermore, we find that clustering of integrins by phase separation of talin is sufficient to activate them even in the absence of ECM. Both our experimental and computational results suggest that interaction force transmission through actin linking enables stabilization and promotes biocondensation of talin. These results demonstrate that biocondensation of talin is sufficient for adhesion nucleation and offer an explanation for the differential formation of IACs around the cell periphery.

## Results

### Optogenetically induced talin-talin interactions through IDRs rapidly and reversibly promote IAC growth and enhances integrin recruitment

While talin is known to be a critical player in adhesion formation, it remains unclear how it might go from activating single integrins to nucleating the formation of new adhesions. To better understand this, we developed optogenetic tools to manipulate talin’s localization and self-association to determine the requirements for talin to nucleate IACs *in vivo*. We reasoned that in addition to recruiting talin to the plasma membrane to activate it and bring it into proximity with integrins, clustering or phase separation of talin might be required to aggregate integrins and form IACs. When combined with its binding partner CIBN, C2 can perform both of these functions simultaneously. By attaching an IDR to C2, such as from the RNA-binding protein FUS, light-inducible phase separation can be achieved. While with C2 alone, primarily membrane localization is observable, but visible membrane droplets are formed across the cell when the IDR is included (**Fig 1A**). These membrane droplets bear more resemblance to the way focal adhesions are distributed across the membrane, although they are more rounded and uniform in morphology.

**Figure 1:**
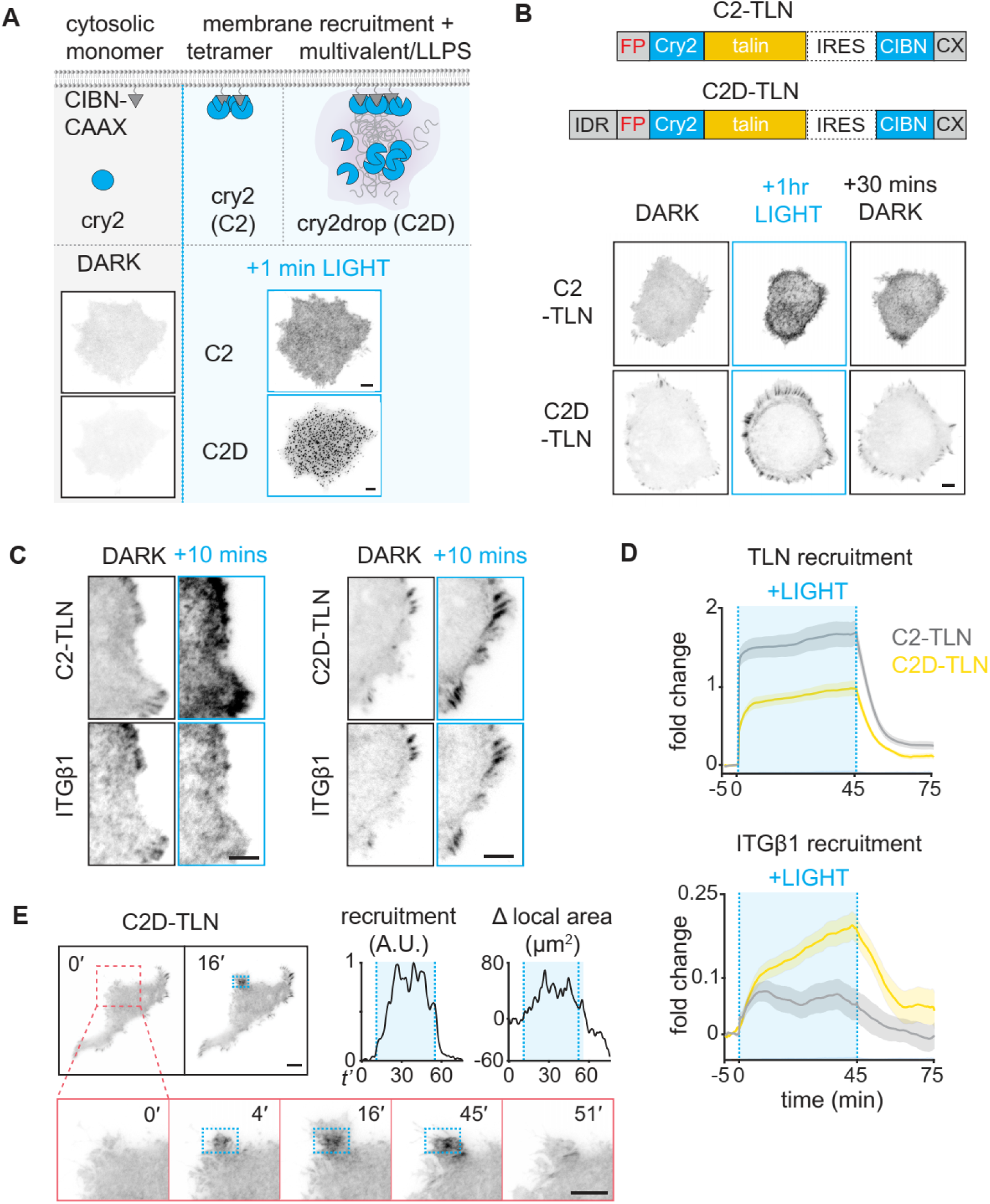
Light-mediated talin self-association drives focal adhesion assembly. **A)** (Top) schematic depicting optogenetic tetramerization of a cytosolic Cry2 (C2) and a membrane CIBN-CAAX under blue light, without (left) and with (right) an IDR fused to Cry2 (C2D). Both methods drive the construct to the membrane and cause self-association when activated, but only with the IDR attached are LLPS droplets formed. (Bottom) TIRF images of HEK 293T cells expressing either of the constructs above before and after one minute of light illumination. **B)** (Top) two optogenetic talin constructs based on the same design as panel A with an IRES between the pieces, one fused with a FUS-IDR (C2D-TLN) and another without it (C2-TLN). (Bottom) representative images of HEK 293T cells expressing snap-C2-TLN (top; n=27) or FusionRed-C2D-TLN (bottom; n=14) immediately before one hour of light activation over the entire cell and after 30 minutes of recovery in the dark. **C**) Cells co-expressing ITGβ1-miRFP680 and either FusionRed-C2-TLN or FusionRed-C2D-TLN before and after light activation over the entire cell and **D)** quantification (n=18 cells each condition, mean ± S.E.M). **E)** (Left) spot activation of C2D-TLN in the blue outlined region and the subsequent recovery. (Right) Quantification of recruitment and protrusion/and retraction in the zoomed area. Scale bars: 5μm; line plots.

When talin is properly oriented and extended, the N-terminus will be membrane-proximal. We therefore chose the N-terminus of talin for fusion to enable C2 binding to CIBN on the membrane not to disrupt this conformation. We fused talin’s N-terminus with either C2 alone or with the IDR from the N-terminus of the RNA-binding protein FUS (C2D), along with either a FusionRed or SNAP fluorescent tag for visualization. We then tested these constructs’ ability to be recruited and reversibly produce IACs in HEK 293T cells plated on fibronectin (FN) by total internal reflection fluorescence microscopy (TIRFM; **Fig. 1B**). HEK 293T cells produce relatively few adhesions, which are located almost exclusively around the periphery where myosin-associated actin is present and retrograde flow occurs. By contrast, the center of the cell contains, little if any, myosin II and ventral actin (**Fig. S1**), thus providing a blank canvas for *de novo* adhesion creation. Within seconds of blue light exposure, both constructs were able to recruit talin to the ventral surface. However, C2D-TLN clustered more efficiently and formed adhesions along the cell periphery, which showed a nearly full recovery after about 20 minutes after ceasing light activation (**Fig. 1B; Movie S1&S2**). To determine if integrins were also recruited to adhesion sites, we observed the recruitment and localization of β1-integrins with a fluorescent tag. Although in the absence of FUS-IDR more talin was localized to the membrane in response to light, integrins accumulated in greater numbers with C2D-TLN and showed more clustered localization (**Fig. 1C, D**). We then attempted to induce local formation of adhesions with FUS-CRY2-Talin through spot illumination using a digital micromirror device (DMD) to pattern blue LED light. Local illumination around the periphery of the cell led to the formation of IACs and a small amount of protrusion, which withdrew when the light was removed (**Fig. 1E; Movie S3**). These results indicate that weak, multivalent interactions between talins, such as those provided by FUS-IDR can help facilitate integrin clustering, IAC formation, and growth.

### Paxillin rapidly complexes with talin in a traction force-independent manner to enable LLPS and FA nucleation

While our optogenetic talin with the FUS-IDR was able to effectively produce adhesions around the edge of cells, neither of the optogenetic constructs produced significant IAC nucleation in the interior regions of the cell. If FUS-IDR were sufficient to drive phase separation of talin on its own, we would expect talin droplets to form across the membrane of the entire cell. Indeed, when only the C2D construct is exposed to light without the talin insertion, FUS-IDR droplets are formed across the cell membrane **(Fig. 2A, top; Movie S4**). We wondered if paxillin, an adaptor protein that binds to talin, might mediate nucleation. Paxillin is often associated with nascent adhesion formation^30^ and has been shown to undergo phase separation *in vitro*^17^. Notably, HEK 293T cells have very little expression of paxillin or its isoform, leupaxin^31^ (**Fig. S2A**). When we co-expressed paxillin with C2D-TLN and exposed cells to blue light, we observed the formation of clusters in the central area of the cell within seconds (**Fig 2A, middle; Movie S5)**. On FN, these clusters are enhanced at the edge, but without ECM, when cells are plated on poly-L-lysine (PL), clusters remain rounded and form more uniformly across the cell (**Fig 2A, bottom; Movie S6**). The initial clusters strongly resemble the LLPS droplets produced by FUS-IDR alone, suggesting that paxillin enables the phase separation of talin. These droplets show hallmarks of LLPS, including fusion and fission **(Fig. S2B; Movie S7**). Notably, paxillin almost directly mirrors the accumulation of talin with very little, if any, time delay **(Fig S2C)**. This, combined with the observation that recruitment does not require ECM, suggests that paxillin can be recruited to talin in the absence of traction force. Previous studies have reported both mechanosensitivity^32^ and mechanoinsensitivity^33,34^ of paxillin recruitment, while this result suggests that there is a significant fraction of paxillin that can be recruited independently of force.

**Figure 2:**
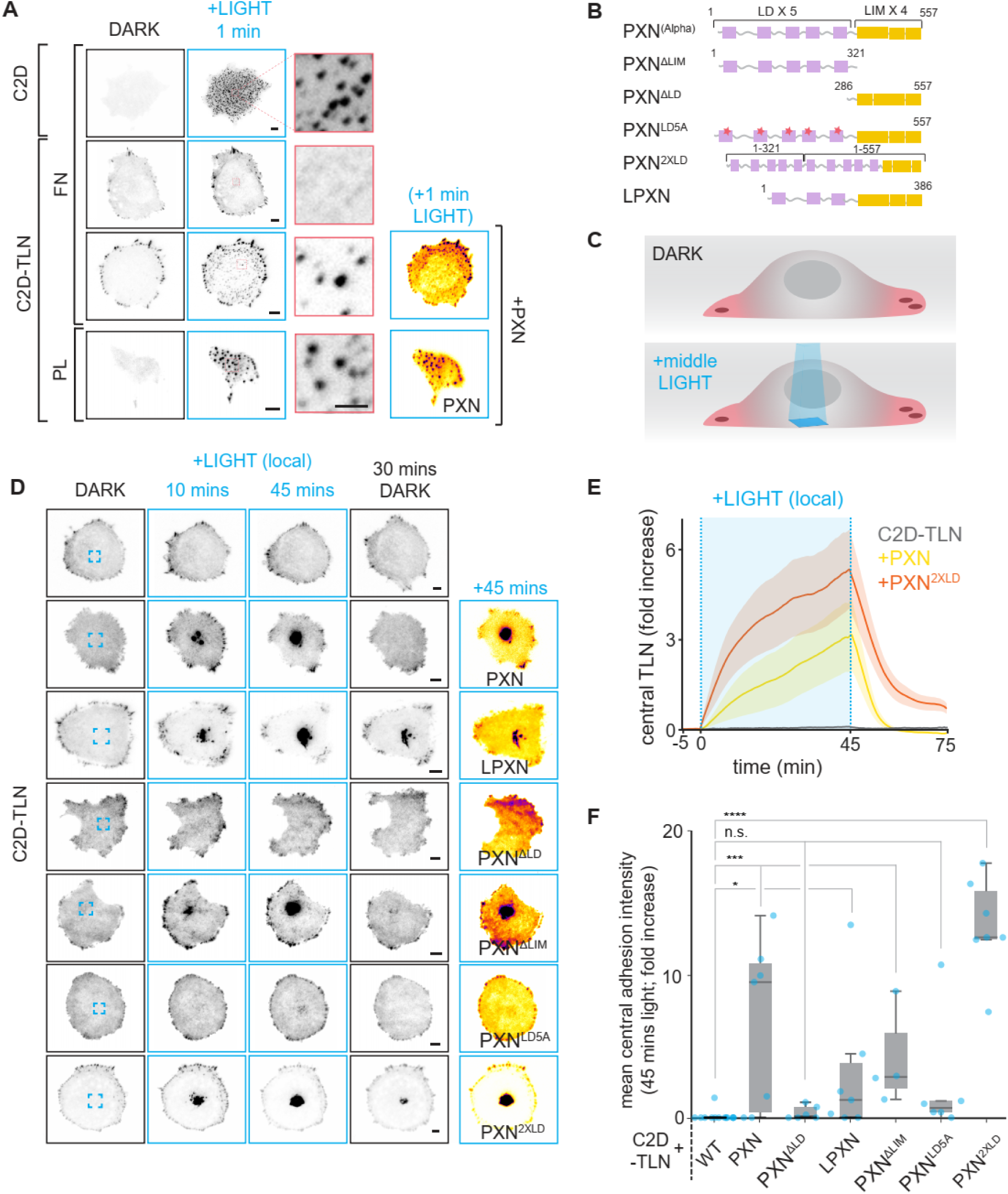
Paxillin enables LLPS of talin and central focal adhesion nucleation. **A)** Representative images of HEK 293T cells before and after 1 minute of blue light exposure expressing FusionRed tagged C2D, C2D-TLN, or C2D-TLN along with PXN-miRFP680 on the coated substrate as indicated (n=>7 cells each). Zoomed regions show the rapid formation of talin droplets. **B)** Depiction of the modified paxillin constructs. **C)** Schematic of central illumination of cells by applying a 12 μm^2^ square region of light to the center of the cell as to activate only that region. **D)** Montage of central activation of FusionRed tagged – C2D-TLN for 10 and 45 minutes, followed by a 30 minute dark recovery with the indicated conditions. **E)** Quantification of the C2D-TLN central recruitment alone, and with PXN-miRFP680 or PXN^2XLD^-miRFP680 (n=>7 cells each, mean ± S.E.M.) during the timecourse in D. **F**) Box and whisker plot showing quantification of the change in central recruitment of C2D-TLN at the 45 minute timepoint (two-sample t-test, n=13,7,6,7,4,6,7 cells, \**P*<0.05, \*\*\**P*<0.001, \*\*\*\**P*<0.0001). Scale bars: 5μm, insets 2 μm.

To understand which regions of paxillin enabled the formation of central talin droplets, we co-expressed C2D-TLN with various truncations and mutations in paxillin (**Fig 2B**). By illuminating only the central regions of cells with blue light (**Fig 2B**) and observing changes in existing IACs and the formation of new talin clusters in the cell center, we assessed the contribution of different domains of paxillin to talin LLPS and recruitment (**Figs. 2D-F**). Consistent with our global activation results, C2D-TLN could not form central IACs alone. However, when additional paxillin or its homologous protein leupaxin was co-expressed, central clusters formed readily. The phase separation functions of paxillin are primarily thought to be mediated by its N-terminus. Paxillin’s N-terminus contains several IDRs as well as binding sites for FAK in its LD domains which have also been shown to mediate LLPS *in vitro*.^16,17^ Consistent with the hypothesis that LLPS was necessary for nucleation of IACs, deletion of the N-terminal IDR and LD domains ablated the ability to form central IACs, while deletion of the C-terminal LIM domains did not prevent central IAC formation. It was previously shown that mutation of an Asp residue in each LD domain which mediate FAK binding to paxillin^35^ to Alanine greatly reduced the phase separation ability of FAK *in vitro*.^16^ When we expressed this mutant version of paxillin (PXN^LD5A^), central IAC formation was also greatly reduced, suggesting that FAK may be important for paxillin-induced LLPS and IAC formation. Consequently, increasing paxillin valency by doubling the LD domains resulted in even more robust recruitment to the center. These results suggest that paxillin’s LD domains enable phase separation of talin, enabling it to form IACs.

### LLPS of Talin enables IAC nucleation

Is the ability to nucleate adhesions a function of talin LLPS or of paxillin? To determine this, we attempted to modify talin to undergo light-induced LLPS without overexpression of paxillin. While talin may participate in LLPS mediated by adaptor proteins, it is possible that talin in an unfolded state could undergo biocondensation independently, as an *in vitro* study has recently shown.^15^ Thus, we modified our C2D-TLN with mutations in the inhibitory interface (M319A, T1767L, and E1770K; C2D-TLN^3CA^), which have been previously shown to disrupt talin’s autoinhibition.^36,37^ Alongside these mutations, we also produced versions of our optogenetic constructs where we replaced the FUS-IDR with the N-terminus of paxillin, including either the first three (C2-TLN^3LDPXN^) or all five LD domains (C2-TLN^5LDPXN^; **Fig. 3A**). Since paxillin normally interacts with integrins,^38^ kindlin,^39^ and possibly the talin head^40^ via its LIM domains, we reasoned that fusing the LD domains to the N-terminus of our optogenetic construct after the fluorophore and C2 might provide similar membrane proximity to natural paxillin. We then tested each construct’s ability to form central adhesions in response to local light activation (**Figs. 3B-D & Movie S8**). All constructs were able to form central IACs in response to local light activation, though the single activating mutation in talin (M319A) had a modest effect (**Fig. 3SA**). C2-TLN^3LDPXN^ and C2D-TLN^3CA^ showed similar levels of IAC formation, while C2-TLN^5LDPXN^ had the most dramatic effect, forming large aggregates which appeared to partition into multiple phases with continued activation (**Fig. S3B).** In cells with C2D-TLN^3CA^, C2-TLN^3LDPXN^, and particularly C2-TLN^5LDPXN^, central IACs sometimes outcompeted the preexisting peripheral adhesions, diminishing them, rather than enhancing them in response to central activation, showing competition between adhesions.

**Figure 3:**
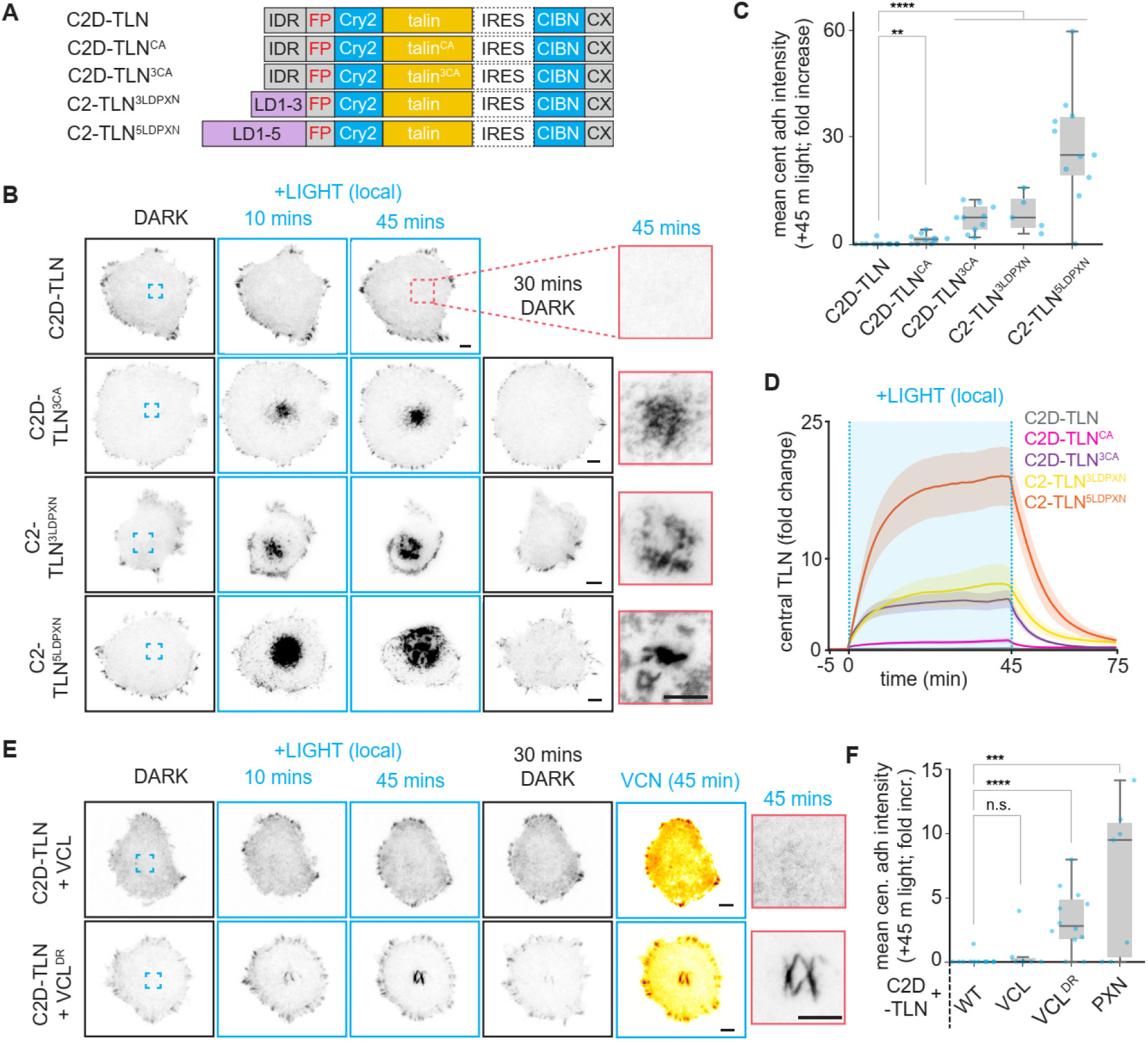
LLPS of talin allows for adhesion nucleation. **A)** Schematic representation of optogenetic constructs for wildtype talin and its variants. **B)** HEK 293T cells with the indicated conditions subjected to 45 minutes of central activation in the region shown in blue. Activated regions after 45 minutes (rescaled for contrast) are shown to the right and **C, D)** quantification (two-sample t-test, n=13,10,10,5,11 cells). **E)** Central activation of C2D-TLN co-expressed with either wildtype vinculin or deregulated vinculin. Activated regions after 45 minutes are shown to the right. **F)** Quantification of the C2D-TLN recruitment in E (two-sample t-test, n=13,7,12,7 cells). Scale bars: 5μm; line plots mean ± S.E.M; \*\**P*<0.01, \*\*\**P*<0.001, \*\*\*\**P*<0.0001).

In previous *in vitro* experiments, talin underwent phase separation when the active conformation was stabilized by a deregulated vinculin (VCL^DR^) in which the residues that mediate autoinhibition had been disrupted (N773A and E775A).^15,41^ We hypothesized that expression of this active vinculin could also enable central IAC formation. Thus, we expressed wild-type vinculin or the deregulated vinculin in our C2D-TLN cells and performed light activation in the center of the cells. Consistent with the idea that phase separation of talin enables the nucleation of IACs, cells co-expressing deregulated vinculin were able to recruit talin to the center of cells, while those expressing wild-type vinculin were not (**Figs. 3E & 3F**). However, central IACs produced in the presence of deregulated vinculin had a more elongated morphology, indicating that increased vinculin activity may be necessary for effective linking to the cytoskeleton. Taken together, these data suggests that LLPS of talin is an effective mechanism for adhesion nucleation.

### LLPS of talin partitions and activates integrins independently of ECM

Since central clusters nucleated by light-induced LLPS of talin often differed in morphology from natural focal adhesions, we sought to determine if integrins were recruited to these clusters. We expressed miRFP680-tagged β1 integrin along with either C2D-TLN alone or in the presence of paxillin overexpression and then applied light to the center of cells. Without paxillin overexpression or other modifications enabling phase separation, the optogenetic tool was not visibly recruited to the center of cells, and consequently, neither was β1 integrin. Conversely, when paxillin was overexpressed, C2D-TLN and integrins were both recruited to the center of cells. We then repeated this experiment using the construct most effectively recruited to the center in previous experiments, C2-TLN^5LDPXN^, instead of overexpressing paxillin directly. This construct was able to recruit integrins as well, which partitioned in concert with the optogenetic tool (**Fig. 4A-C; Movie S10**). Furthermore, integrins were often depleted from the peripheral adhesions as they were recruited centrally, revealing competition between adhesions. Conversely, FUS-IDR clustering in the absence of talin forms droplets but does not recruit integrins (**Fig S4**).

**Figure 4:**
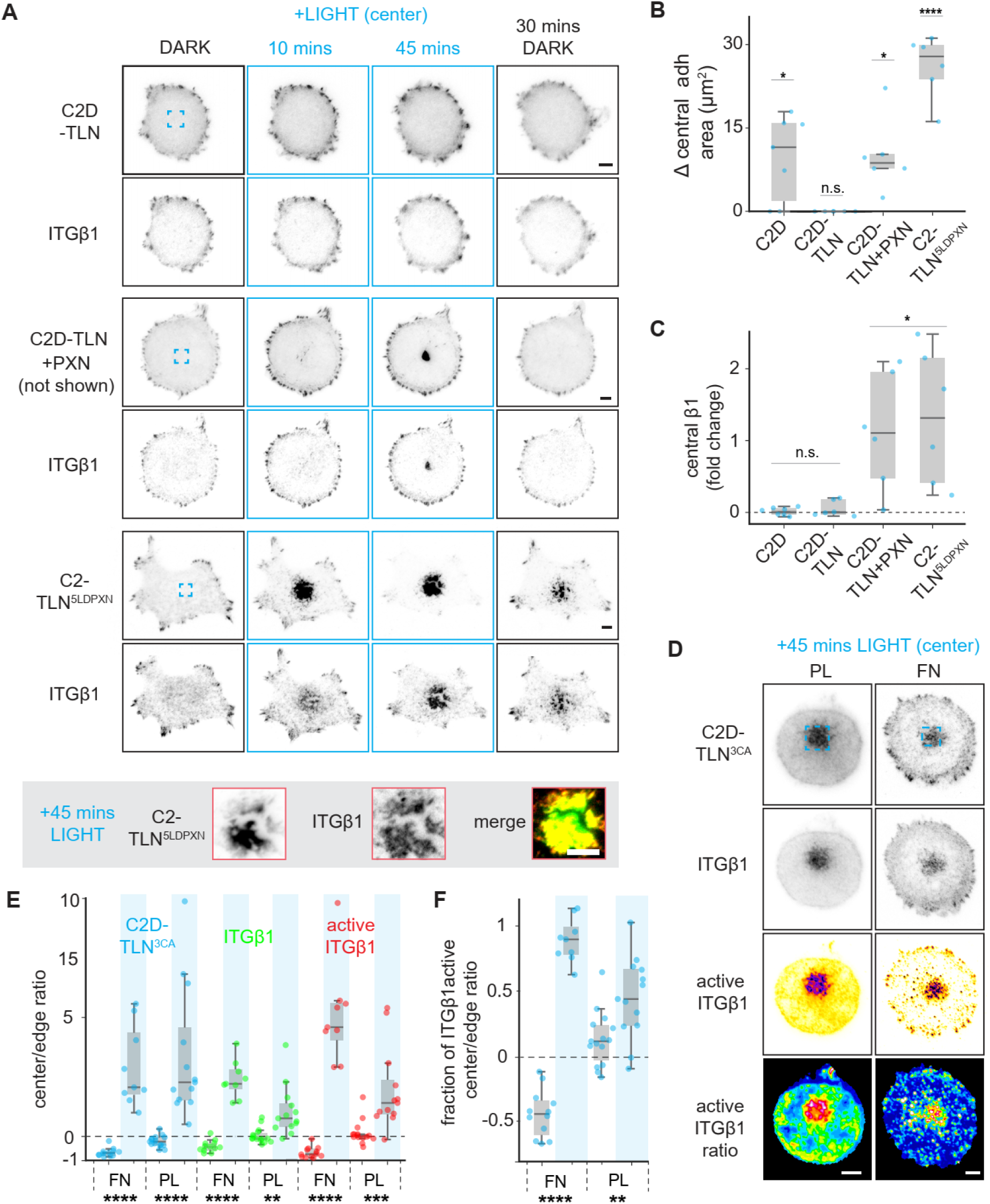
Phase separation of talin partitions and activates integrins. **A)** Montage of HEK 293T cells co-expressing (top) ITGβ1-miRFP680 and FusionRed-C2D-TLN and (middle) with EGFP-PXN, or (bottom) expressing C2-TLN^5LDPXN^ instead of C2D-TLN exposed to central illumination in the blue box for 45 minutes before being allowed to recover for another 30 mins. Rescaled zooms of the central activated regions at 45 minutes are shown below the panel. **B)** Quantification of the change in central adhesion area or change in **C)** central β1integrin after 45 minutes of activation (one-sample t-test, μ_0_=0, n=>6 cells each condition). **D)**. Representative images of HEK 293T cells co-expressing FusionRed-C2-TLN^3CA^ and ITGβ1-miRFP680 plated on either fibronectin or polylysine and fixed after 45 minutes of central activation. Active ITGβ1 was visualized by conformation specific antibody (9EG7). The ratio of active-to-total integrin is shown to the right. **E)** Intensity quantification of the cell center-to-edge intensity ratio of each construct and active integrin staining with and without 45 minutes central activation (blue highlighted bars). A ratio of zero indicates that the center and edge have equal amounts, a ratio of one would indicate a center that is enriched by 100%, and conversely, −1 would indicate the edge enriched by 100% relative to the center. **F)** Center/edge ratio of the fraction of total integrins which are active with (blue highlights) and without 45 minutes light stimulation on either PL or FN (two-sample t-test between the light and dark of each condition, n=>9 cells each condition; \*\**P*<0.01, \*\*\**P*<0.001, \*\*\*\**P*<0.0001). Scale bars: 5μm.

To determine whether these integrins are maintained in the activated conformation in the phase-separated central clusters we fixed cells containing both C2D-TLN^3CA^ and ITGβ1-miRFP680 after 45 minutes of central activation and stained for active β1 integrin (9EG7; **Fig. 4D**). To compare the relative levels of activation we calculated the ratios of the center intensities with those around the periphery (**Fig. 4E)**. We subtracted one from this ratio, such that equal amounts in both regions would be equal to zero. With natural adhesion formation on FN, we would expect this ratio to be negative in unstimulated cells on FN and zero on PL. Indeed, this was the case for all three markers when cells were unstimulated. However, in cells which received central optogenetic stimulation all three markers were increased significantly in the center on both FN and PL. Central optogenetic recruitment of C2D-TLN^3CA^ was similar on both PL and FN, while β1 integrin recruitment and active β1 were more present after activation on FN, although still enriched on PL. Strikingly, when we ratioed the relative amount of active β1 integrins with the total, the fraction of central integrins active after stimulation remained elevated by more than 40% even in the absence of ECM (**Fig. 4F**). These results suggest that the inside-out signal triggered by light-induced LLPS of talin can be sufficient for integrin activation.

### Talin condensates nucleate actin and are stabilized by peripheral actomyosin

As optogenetically induced IACs were able to recruit and activate integrins, we sought to understand if they also engaged with actin. To determine this, we visualized actin via Lifeact-miRFP680 while optogenetically activating C2-TLN^5LDPXN^ in the center of cells. Prior to light activation, most cells displayed little central ventral actin. However, as central light activation persisted, most cells slowly began to accumulate actin around C2-TLN^5LDPXN^ droplets (**Figs. 5A & 5B; Movie S11**). This actin typically persisted longer than the optogenetic construct after the light was removed, though it was still significantly less than what was found around the periphery of the cell and differed in morphology. Thus, while central IACs were able to engage some actin, it was distinct in from those adhesions around the periphery with access to actomyosin and the contractile force associated with it.

**Figure 5:**
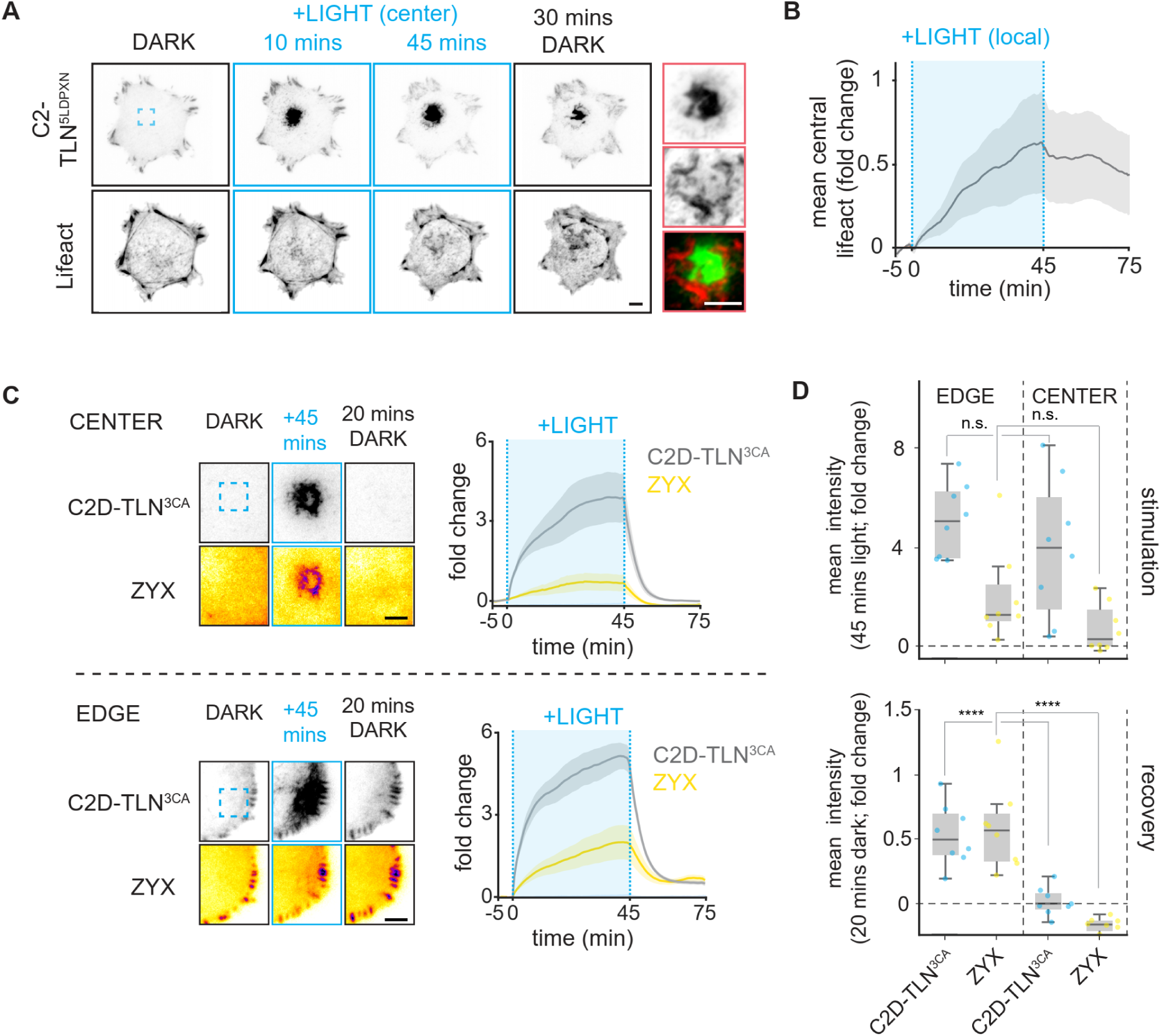
Actin engagement stabilizes talin recruitment. **A)** Montage of HEK 293T cells co-expressing Lifeact-miRFP680 and FusionRed-C2-TLN^5LDPXN^ exposed to central illumination in the blue box for 45 minutes before being allowed to recover for another 30 mins. (Right) zoom of activated regions and merged channels with Lifeact shown in red and C2D-TLN shown in green. **B)** Quantification of central Lifeact accumulation in A (n=18 cells; mean ± S.E.M.). **C)** Spot activated regions (top panel: center activation; bottom panel: edge activation) of cells expressing FusionRed-C2D-TLN^3CA^ and ZYX-miRFP680 after 45 minutes stimulation and 20 minutes recovery. **D)** Quantification of fold changes and mean intensities (two-sample t-test, n=8 cells each condition, \*\*\*\**P*<0.0001). Scale bars: 5μm.

Based on our observations, we hypothesized that in the absence of light adhesions are stabilized by binding to actin and force transduction. Since ventral actin and myosin II are primarily located around the edge of the cell (**Fig. S1**), we reasoned that talin recruited there might be more stable than those recruited at the center. Thus we compared the recruitment at the edge of the cell with the central regions for C2D-TLN^3CA^ and zyxin, an *α*-actinin binding^42^ focal adhesion protein that is responsive to force^43,44^, (**Fig. 5C & 5D, Movie S12**). Optogenetic C2D-TLN^3CA^ was recruited at comparable levels in both regions of the cell. However, while zyxin was co-recruited to the edge in all cells, it was recruited to the center in only half of the cells. After the light was removed, we observed the reversion of both constructs to determine if force and actin engagement at the edge would affect the constructs’ recovery. After 20 minutes in the dark, both C2D-TLN^3CA^ and zyxin remained elevated at the edge of the cell by about 50% relative to their initial levels. Conversely, there was no elevation of either construct in the center of the cell after the same recovery time. These data are congruent with the hypothesis that forces around the periphery of cells can stabilize integrin activity even after optogenetic stimulation has ceased, while a lack of force stabilization in the middle of the cell prevents adhesions from forming naturally in this region.

### A 2D-lattice model of LLPS-induced IAC formation

To further understand the interplay between phase separation and mechanotransduction processes mediated by talin in regulating the size of focal adhesions at the cell edge and the nucleation of biocondensates in the cell center, we developed a model of molecular clutch formation (**Fig. 6A**) based on a recently published study^45^. The model considers the formation of molecular clutches between the actin cytoskeleton and the substrate by integrin, talin, and vinculin. This process is guided by phase separation of talin molecules that can be induced either through unfolding of the mechanosensitive R3 domain of talin^15^ or via interaction of the IDRs on optogenetic constructs expressed in cells. To this aim, molecular clutches were modeled as individual elements on a 2D grid representing the interface between a cell and the underlying substrate. These elements participate in neighbor-neighbor interactions that promote the formation and stabilization of molecular clutches in adjacent sites. The model also takes into account mechanotransduction processes, such as force-induced dissociation of molecular clutches and unfolding of the mechanosensitive R3 domain of talin. Unfolding of the R3 domain enables interaction with vinculin, which stabilizes molecular clutches and promotes focal adhesion growth in response to mechanical load^46^.

**Figure 6:**
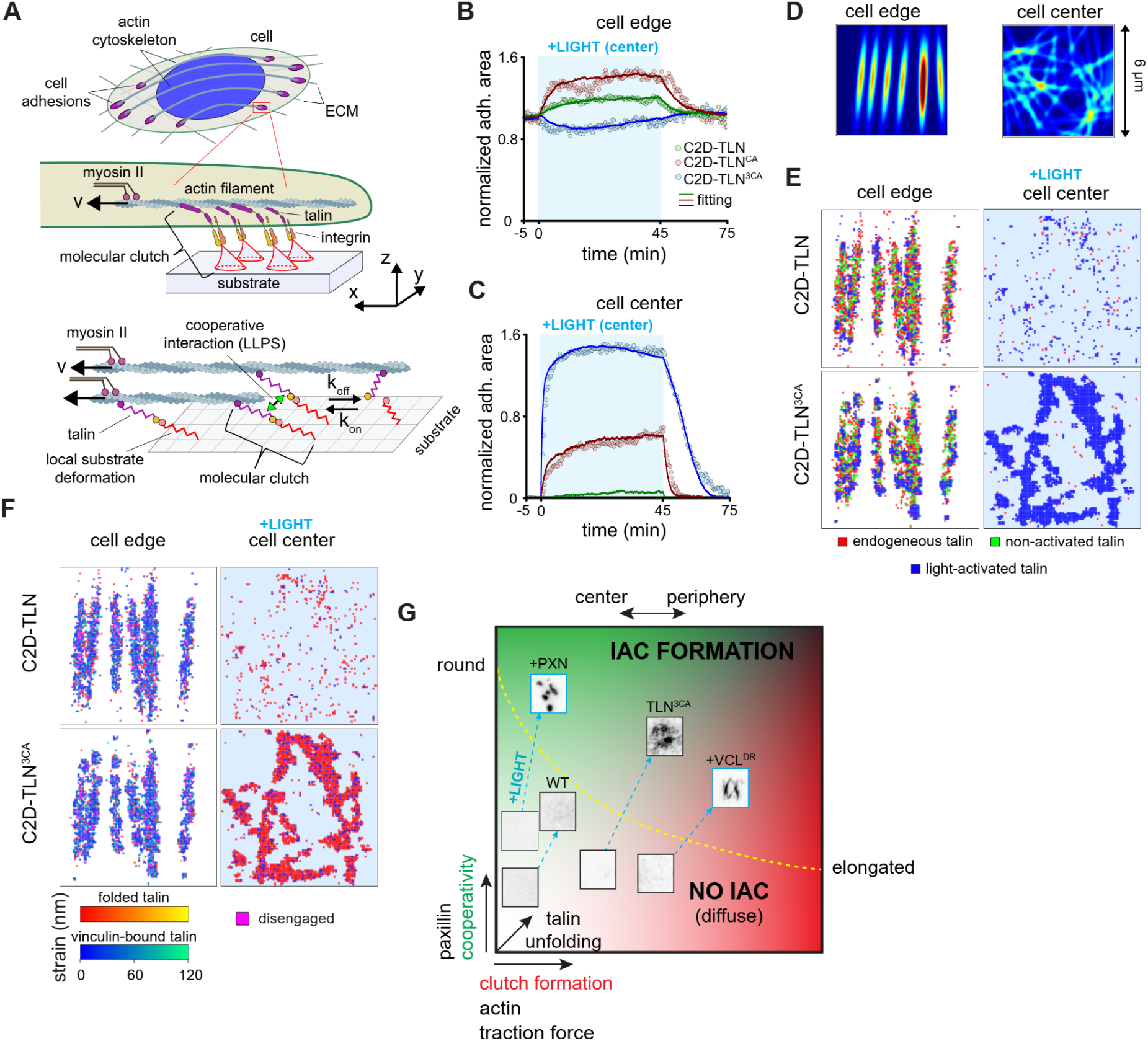
Model of adhesion nucleation by LLPS. **A)** Top panel: integrins and adaptor proteins such as talin are the main components of cell adhesion linkages that form between the actin cytoskeleton and substrate. Myosin II-generated retrograde flow of actin filaments (v) leads to mechanical stretching of these links, causing local substrate deformations (shown in red). Force-transmitting units, each including a cellular part (integrin and adaptor proteins such as talin) and an extracellular part (locally deformed substrate), are referred to in this study as ‘molecular clutches’. Bottom panel: in the model, molecular clutches are represented by composite springs consisting of two parts – an intracellular one and an extracellular one. The substrate in the model is described as 2D grid, at the nodes of which molecular clutches can form. The kinetics of molecular clutches under mechanical load generated by myosin II motors is described by their formation and dissociate rates, k_on_ and k_off_,45 respectively. In addition to mechanical loading, the values of these rates are affected by the type and the state of neighbouring molecular clutches, which can undergo cooperative interactions due to LLPS between different types of talin molecules used in the experiment. **B & C)** Normalized total projected area of IACs at the cell edge (B) and in the cell center (C) as a function of time. Solid lines show experimental data fitting obtained by averaging over three independent stochastic simulation of cell adhesion complexes. **D)** Representative image of the density function of sites available for the formation of molecular clutches used in the model to represent focal adhesions at the cell edge and a randomly scattered filamentous network in the cell center. **E & F)** Representative images of the light-activated state of molecular clutches (E) and their type (talin-folded, talin-unfolded, vinculin-bound), distribution and mechanical strain (F) at the edge and in the center of cells expressing C2D-TLN and C2D-TLN^3CA^ talin constructs after their light activation. The results were obtained by performing stochastic simulations based on the model shown in panel (A). **G)** Schematic phase diagram showing the regimes in which IACs can form. At the cell edge, clutch formation (red) is higher, allowing for a lower cooperativity (green) threshold to form adhesions, while in the center cooperativity must be very high to nucleate IACs. Paxillin availability increases cooperativity, while talin unfolding increases both cooperativity and clutch formation. Optogenetic activation increases both cooperativity as well as clutch formation to a lesser extent, allowing crossing of the phase boundary where IAC formation can occur.

The model was then used to fit experimentally measured kinetics of time-dependent size change of IACs both at the center and periphery of the cell upon center light-induced activation of optogenetic talin constructs (**Figs. 6B & C**). Both the model and experimental data showed that there are two distinct regimes of talin recruitment in response to central light activation which dependent on the cooperativity and clutch formation rate (**Figs. 6D-F; Movies S13 & S14)**. The first regime is characterized by an increase in clutch formation at the cell edge but little change in the middle of the cell upon central light activation, as observed for the C2D-TLN. Fitting the experimental data to the model showed that this regime is valid for a cooperative binding energy of optogenetic talin constructs with less than ∼2 k_B_T (this energy corresponds to the change in free energy of a molecular clutch containing a light-activated optogenetic talin construct surrounded by other molecular clutches of a similar type). This is consistent with the ∼1.4 k_B_T we measured for C2D alone in our system (**Fig. S5A**).

The second regime, under high cooperativity (greater than ∼4 k_B_T), showed strong nucleation and phase separation potential, such as with C2D-TLN^3CA^ and C2-TLN^5LDPXN^. However, in both cases IACs could form at the cell edge, even prior to light activation. This is because clutch formation fixes talin in place and under force leads to talin unfolding. This both increases cooperativity and lowers the threshold for IAC formation. Fitting of the experimental data also revealed that phase separation of the active talin constructs should result in an increase in the formation rate of molecular clutches in the light activated region due to a local increase in the concentration of phase separated talin constructs. Thus, clutch formation and cooperativity synergically enable biocondensation of talin and IAC formation **(Fig. 6G)**. These results suggest that the processes responsible for regulating the phase separation behaviour of molecular clutch components, such as talin state, or the availability of other adaptors, play an important role in governing the competition between adhesion complexes, differentially altering the stability and adhesion nucleation potential

## Methods

### Plasmid construction

All plasmids were generated via seamless cloning using the ClonExpress Ultra One Step Cloning Kit (Vazyme, C115/C116). Amplification primers containing recombination sequences were designed according to the manufacturer’s protocol. Inserts were amplified using either Phanta Flash Super-Fidelity DNA Polymerase (Vazyme, P510) or Phanta UniFi DNA Polymerase (Vazyme, P516), following the manufacturer’s protocol.

The C2D construct was created by subcloning the FUS-FusionRed-Cry2PHR^29^, along with IRES-CIBN-CAAX2 into the pBJ (piggyBac transposon) vector (Addgene #176531). The C2 construct was generated by removing FUS from C2D via PCR. Subsequently, C2 and C2D were linearized by restriction enzyme digestion and ligated with full-length human talin-1^47^ to generate C2-TLN and C2D-TLN, respectively. The constructs C2-TLN^3LDPXN^ and C2-TLN^5LDPXN^ were generated by replacing the FUS-IDR in C2D-TLN with the N-terminus from pmCherry Paxillin (Addgene #50526; residues 1–215 for 3LD, residues 1–299 for 5LD). Point mutations were introduced to produce active talin variants^36,37^ C2D-TLN^CA^ (M319A), C2D-TLN^3CA^ (M319A/T1767L/E1770K), and VCL^DR^ (N773A, E775A)^41^. MiRFP680-tagged full-length wild-type paxillin, its truncations or point mutations, along with integrin β1, zyxin, and Lifeact, was achieved by subcloning into the pcDNA3.1 backbone. Plasmid sources and primers used for cloning are listed in **Tables S1&S2**.

### Generation of cell lines and cell culture

HEK-293T^1^ and U2-OS^6^ were cultured in Dulbecco’s Modified Eagle Medium (DMEM, Gibco, 10569044) while PC-3^7^ and DU145 (a generous gift from Dr. Hailiang Hu, Department of Biochemistry, School of Medicine, Southern University of Science and Technology, Shenzhen, China) cells were maintained in Roswell Park Memorial Institute (RPMI)−1640 medium (Gibco, 11875093). All media were supplemented with 10% fetal bovine serum (FBS; Gibco, A5256701) and 1% penicillin-streptomycin (Gibco, 15140122). Cells were incubated at 37°C in a humidified 5% CO₂ atmosphere.

Cells were transfected using Lipofectamine 3000 according to the manufacturer’s instructions. For expression of ITGβ1-miRFP680, cells were transfected at least 48 hours before imaging to reduce the amount of tagged integrins trapped in the endoplasmic reticulum (ER). Stable cell lines for optogenetic tools were generated via the PiggyBac transposon system. Target pBJ plasmids were transfected into HEK-293T cells at pBJ plasmid: transposase molar ratios of 1.2:1. Stable cell lines were then selected with 2 μg/mL puromycin (Thermo Fisher, A1113803) for one week, beginning 24 hours post-transfection.

### Live cell imaging and optogenetic stimulation

Before imaging, 96-well glass-bottom plates (Cellvis, P96-1.5H-N) were coated with poly-L-lysine (PL; Sigma-Aldrich #A-005-C) or fibronectin (FN; Corning, #354008). For PL coating, wells were incubated with 50 μL of 0.01% w/v PL for 30 minutes at room temperature, washed five times with DPBS (Gibco, 14190250), and dried before cell seeding. For FN coating, a 1 mg/mL FN stock solution was prepared by dissolving fibronectin in DPBS and incubating at 37°C for 30 minutes, then aliquoted and stored at −80°C. Each well was coated with 50 μL of 20 μg/mL FN working solution (diluted from stock in DPBS) for 30 minutes at 37°C, washed twice with DPBS, and dried. To minimize cell aggregation, approximately 800 HEK-293T cells were plated per well. Cells were spread for at least 2.5 hrs on PL-coated plates in serum-free DMEM or on FN-coated plates in DMEM with 4% FBS.

Imaging was performed using total internal reflection fluorescence (TIRF) on a Nikon Ti2 inverted microscope with a motorized TIRF module, Photometrics Kinetix back illuminated sCMOS camera and a 60× oil-immersion objective (NA 1.49). Optogenetic illumination was performed at 435 nm using a Mightex Polygon 1000 DMD equipped with a CoolLED pE-800 by pulsing for 60ms at approximately 1 mW/cm^2^ at 20-second intervals. For local activation experiments, DMD illumination was restricted to a 12 μm^2^ square region, in the center or at the edge of the cell, per the experiment. A custom macro was used to change illumination patterns between XY positions to prevent cross-activation.

### Immunofluorescence

For detection of active integrins, stable cell lines co-expressing C2D-TLN^3CA^ and ITGβ1-miRFP680 were activated in central cellular regions for 45 minutes on PL or FN coated plates. Cells were immediately fixed post-activation with 3% paraformaldehyde (prepared from 32% stock; Electron Microscopy Sciences, 15714) in DPBS for 10 minutes at room temperature. Following fixation, cells were washed three times with DPBST (DPBS + 0.05% Tween 20) and blocked overnight at 4°C with a non-permeabilizing solution consisting of 3% BSA and 10% goat serum (Sigma-Aldrich, G9023) in DPBST. After blocking, cells were gently rinsed once with DPBST and incubated with primary antibody against active integrin β1 9EG7 (BD Pharmingen™, 553715; 5 μg/mL) for 3 hours at room temperature. Cells were then washed five times with DPBST and incubated with species-matched secondary antibody (Invitrogen, A-11006), diluted 1:1500, for 1 hour. After five additional washes with DPBST, samples were imaged by TIRFM.

### Total RNA extraction and Real-time PCR

Total RNA was extracted by RNAzol® RT (RN 190, MRC) following the manufacturer’s protocol. cDNA was synthesized from mRNA templates with the HiScript IV cDNA Synthesis Kit (Vazyme, R412). qPCR was performed using a Bio-Rad CFX Opus system with SupRealQ SYBR Master Mix (Vazyme, Q713). Primers are listed in **Table S3**.

### Image analysis

Images were converted to 16-bit TIFF and single cells were cropped and registered in ImageJ2 using the descriptor-based series registration plugin^48^, if necessary. If additional cells were present in frames they were manually removed from the images for analysis. Poorly spread, clumped, or mitotic cells, as well as cells with significant visible ER trapping of integrins, were excluded from analysis. For fixed samples stained for integrin activation, cells which did not still show tool or integrin recruitment after fixing were excluded from analysis. In MATLAB, cells were segmented from the background by thresholding using the non-optogenetic channel if available. The mask is then cleaned up by morphological closing using a 0.5 μmdisk, removal of regions smaller than 7.5 μm^2^, and the filling of interior holes. Using the cell mask images are background subtracted. The first frame of each timelapse is used to calculate the relative expression of that protein by taking an average intensity of the cell. Since LLPS is a concentration-dependent phenomenon, cells which do not have at least 75% of the median expression value of each construct were excluded from analysis, to ensure similar expression across cells. To quantify the fold change in a protein on the inner or outer region of the cell a mask of optogenetic stimulation area is created and dilated by a 1.2 μm disk to account for small drifts and blurring of the area. This area as is further dilated by a 2.4 μm disk to create a buffer and everything which does not fall in this large disk in the cell mask is consider “outside”. The average intensity in each area is then calculated. This intensity is then normalized by average of the dark intensities in the future activation region for local activation experiments. The fold change at a time of activation is calculated using this normalized intensity and subtracting one from it. The fold change in adhesion intensity is done in the same way as cell intensity but using the average intensity of the segmented adhesions in that region rather than the whole cell intensity average in that region. To segment adhesions a 2D 11 μm^2^ square median filter is applied to the image and then thresholded. A single threshold is used for all cells and conditions. Regions of the median filter mask which are not at least 2 standard deviations above the cell mean in intensity are then excluded. Adhesions below 1.5 μm^2^ are removed before morphological opening and closing to clean up the mask. These masks area then used to calculate adhesion area in a region. Protrusion and retraction were calculated as done previously, by calculating the difference between cell masks in each frame.^49,50^ Plots of time courses of optogenetic activation were smoothed over a 7 minute moving average within each stimulation period. For box-and-whisker plots each time point was averaged over 3 points in a one-minute interval. P-values for t-tests are listed in **Table S4.**

### Computational Model

In our model, we aimed to capture the most essential mechanisms underlying formation and maturation of cell adhesion complexes, such as phase-separation processes and force-mediated enhancement of focal adhesions, using a minimal setup in which molecular clutches formed by the proteins, such as integrin, talin and vinculin, were simulated as individual elements on a 2D grid (100×100 sites) representing the interface between a cell and the underlying substrate. The model considers formation of molecular clutches between the actin cytoskeleton and the substrate, which is guided by phase-separation of talin molecules that can be induced either by unfolding of the mechanosensitive R3 domain of talin or via light-induced activation of the optogenetic protein domain (C2) and other phase-separating protein domains attached to talin (FUS, paxillin IDRs). It was assumed that each molecular clutch upon its formation occupies one site in the 2D grid, see **Figure 6A**.

Depending on the type of talin molecules (WT or modified) involved in the formation of molecular clutches and their state (folded / unfolded mechanosensitive R3 domain of talin, light-activated C2 domain or not), the molecular clutches occupying 2D grid were assumed to be in one of the following states:

1) Molecular clutch containing wildtype talin (endogenous, non-modified talin) whose mechnosensitive R3 domain is in the folded state (i.e., folded WT talin).
2) Molecular clutch containing wildtype talin whose mechnosensitive R3 domain is in the unfolded state (i.e., unfolded WT talin).
3) Molecular clutch containing unfolded WT talin bound to vinculin.
4)-6) Same as 1)-3), only in this case the molecular clutch contains an exogenously expressed talin construct with optogenetic domains fused to it (C2, FUS, etc.). The C2 domain of the talin construct is in the inactivated state.
7)-9) Same as 4)-6), only in this case the C2 domain of the talin construct is in the light-activated state.
10-16) Same as 3-9), only in this case the integrin-ECM bond is ruptured in the molecular clutch (the molecular clutch is disengaged from the substrate).

The dynamic behaviour of the system was described in the model by considering stochastic transitions between the above states in much the same way as in a recently published study^45^, taking into account the following kinetic processes:

1) Formation of molecular clutches at the nodes of 2D grid, described by kinetic rate k_on_. It is assumed that upon formation, talin in molecular clutches is in a mechanically relaxed folded state.
2) Force-induced unfolding of the mechanosensitive R3 domain of talin in molecular clutches, caused by their gradual stretching due to the retrograde actin flow generated by the contractile activity of myosin II motors. This process was described by the rate k_u_(F), which depends on the force F applied to the molecular clutch.
3) Refolding of the mechanosensitive R3 domain of talin in molecular clutches, described by the rate k_f_(F), which is also sensitive to the mechanical load applied to the molecular clutch.
4) Vinculin binding to and dissociation from the unfolded mechanosensitive R3 domain of talin in molecular clutches, which were described by the rates k^R3v^_on_ and k^R3v^_off_, respectively. Binding of vinculin to the mechanically unfolded talin in molecular clutches is known to result in force-dependent enhancement of focal adhesions at the cell edge, which was represented in the model by an increased value of the rate of molecular clutch formation, k_on_, at the 2D grid nodes adjacent to molecular clutches with vinculin-bound talins.
5) Rupture of the bond between integrin and substrate in molecular clutches, which is described by the previously experimentally measured integrin-fibronectin bond dissociation rate, k_off_(F), which depends on the molecular clutch tension, F. Rupture of this bond results in a disruption of the force transmission between the actin cytoskeleton and the substrate. However, this does not lead to immediate complete disassembly of molecular clutches in the model as soon as their talin molecules are bound to vinculin (stabilizes molecular clutches) or are involved in interaction with neighbouring molecular clutches, which is mediated by phase-separating domains of talin constructs used in the experiment (C2, FUS, paxillin IDR).
6) Re-establishment of molecular clutches that were previously disengaged from the substrate, resulting in the restoration of the force-transmitting link between the actin cytoskeleton and the substrate, – a process, which is described by the rate k^0^.
7) Complete dissociation of molecular clutches disengaged from the substrate due to dissociation of vinculin from their talin molecules (process # 4 above) or due to rupture of the bonds mediating phase-separation of talins in adjacent molecular clutches containing modified talin constructs (between C2 / FUS / paxillin IDR domains). The kinetic rate that describes the latter process, k_off,t_(ε), is a function of the cooperative binding energy of talins, ε, which determines the strength of their phase-separation (see below).
8) Finally, the deactivation of the light-induced optogenetic domain C2 of the talin constructs was described using the rate k_light-off_ = 0.011 s^−1^, measured in Shin et al., 2017^29^.

A detailed description of most of the above kinetic rates, their values, and their susceptibility to mechanical load applied to molecular clutches by the actin cytoskeleton can be found in Liu et al., 2025^45^. There are only three major differences between this study and that one:

1) It was assumed that not only molecular clutches with vinculin-bound talin (force-induced enhancement of focal adhesions), but also those containing talin constructs with optogenetic domains, affect the formation rate of molecular clutches at adjacent sites of the 2D grid representing the substrate. For simplicity of the model, it was assumed that only molecular clutches containing talin molecules with the light-activated C2 domain induce increased molecular clutch formation at adjacent sites due to LLPS interaction between the optogenetic domains in adjacent molecular clutches:

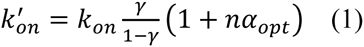

Where k’_on_ and k_on_ are the formation rates of molecular clutches by talin constructs containing the optogenetic domains and endogenous talin, respectively. γ is the fraction of talins containing the optogenetic domain, which diffuse in the cell cytoplasm. n is the total number of sites on the 2D grid (from 0 to 4) occupied by molecular clutches containing talin molecules with light-activated C2 domains that are adjacent to a given unoccupied site at which the formation of a new molecular clutch is considered. α_opt_ is a factor describing enhancement of the formation of a new molecular clutch by adjacent molecular clutches containing talin molecules with the light-activated C2 domain (i.e., LLPS interactions between such molecular clutches and freely diffusing talin molecules effectively result in increased talin concentration at the neighbouring sites).
2) It was assumed that the phase-separating domains of talin constructs (C2, FUS, paxillin IDR) in molecular clutches occupying adjacent sites on the 2D grid, exhibit affinity for each other – the more adjacent molecular clutches contain talin constructs with optogenetic domains, the less likely it is that a given molecular clutch with optogenetic domains will undergo the kinetic process # 7 described above, resulting in complete dissociation of the molecular clutch. This effect was approximated in the model using the following formula:

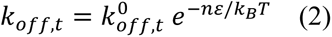

Where k_off,t_ is the dissociation rate of the optogenetic domains of adjacent molecular clutches. ε is the cooperative binding energy of the optogenetic domains. n is the total number of sites on the 2D grid (from 0 to 4) occupied by molecular clutches containing talin molecules with light-activated C2 domains that are adjacent to a given molecular clutch. k_B_ is the Boltzmann constant, T is the temperature.
3) To compare the model predictions with experimental measurements, two general cell regions were considered in the calculations: 1) the cell edge enriched in focal adhesions, and 2) the cell center, which was illuminated with light in experiments to induce optogenetic phase-separation of various talin constructs. To model the spatial heterogeneity of the density of sites available for the formation of molecular clutches at the cell edge and in the cell center, the rate of molecular clutch formation was modulated using the density function, ρ(x): k_on_ = k^0^ ·ρ(x), 0 ≤ ρ(x) ≤ 1, where k^0^ = const is the maximum possible rate of molecular clutch formation. The density function was described by randomly scattered elongated ellipses aligned in nearly the same direction at the cell edge to mimic focal adhesions that typically form in this region; whereas in the case of the cell edge, the density function was described by randomly scattered filaments, see schematic **Figure 6D**.

## Discussion

Optical manipulations of talin, either by disconnecting it at different points in the rod domain^51,52^, by controlling its localization to the plasma membrane^53^, or by controlling talin dimerization^54^ have been demonstrated previously. These tools have been valuable for understanding mechanotransduction through talin. However, these studies mainly focus on talin cleavage and adhesion disruption across the entire cell. To our knowledge, no talin optogenetic tool has previously been used to create IACs, much less in a subcellular manner. Our tools represent a first step towards this goal as they can be used to locally generate IACs with high spatial and temporal precision on a subcellular level. Our observation that C2D-TLN did not form visible central droplets on its own is likely due to the large size of talin-1 (270kDa), compared to the FUS-IDR (20kDa). Furthermore, the optogenetic talin may be dimerized with endogenous talin molecules that do not have a FUS-IDR. For synthetic condensation of talin by FUS-IDR alone, the ratio of FUS-IDR to talin would likely need to be increased significantly.

Talin has been shown to undergo phase separation *in vitro* on the membrane when it is in its unfolded state. However, how could talin achieve the critical concentration to undergo phase separation *in vivo* at an early stage of adhesion formation? LLPS inherently has a lower threshold for occurring on a membrane than in the cytosol^55,56^, and as IACs are made of many proteins, talin does not need to undergo biocondensation alone. Previous studies have shown that numerous other focal adhesion proteins undergo phase separation *in vitro*, many of which may co-phase separate with talin, working synergistically to enable talin to form biocondensates. As such, our results show that paxillin, through its LD domains - which have been shown to undergo LLPS *in vitro* and *in vivo*^16,17^ - greatly enhances the phase-separation and nucleation potential of talin. In addition, we demonstrate that the modulation of paxillin expression level results in an all-or-none response in the formation of central talin clusters – a hallmark of phase separation.

As some other LD domains can bind to talin in its folded form^57,58^, it is not unreasonable that paxillin or other adaptors also bind to talin in the cytosol. Previous work has suggested that focal adhesion proteins, including paxillin can form cytosolic complexes.^23^ Our results show that paxillin was recruited rapidly with talin, with little, if any, delay. This occurred even in the absence of ECM, suggesting that some paxillin may bind talin even in the cytosol. Further evidence that paxillin is rapidly recruited is provided by the observation that paxillin is one of the first proteins recruited to nascent adhesions.^59^ Paxillin is thought to bind to talin in both its R8^58^ and head domains^40^ and with its LD and LIM domains respectively. In addition, it also binds kindlin and the integrin, and thus it may assist in oligomerizing talin dimers further.^39,40^ The dramatic effect generated by attaching the LD domains of paxillin to talin we observed highlights the potential of paxillin to provide this function, even in the absence of the LIM domains. Kindlin, which interacts with paxillin, integrins, and talin^60^, has also been shown to undergo phase separation^14^ and is typically membrane-localized. Furthermore, superresolution studies further confirm that these proteins all lie within the same plane relative to the membrane.^61,62^ Thus, it seems plausible that a small amount of talin recruited to the membrane could cooperatively reach the critical concentration for phase separation with other co-phase-separating proteins, thereby providing the stability needed for nascent adhesion formation.

Previous work has suggested that LLPS plays a role in adhesion nucleation^15–17^, but this hypothesis has not been directly tested *in vivo*. Here, we directly demonstrate how biocondensation of talin could mediate adhesion nucleation. Our results show that phase separation of talin can sequester integrins, activate them, and nucleate IACs. But how does phase separation of talin achieve this? A recent *in vitro* study^63^ showed that ligand, but not talin alone, was able to induce switching to the extended conformation. In contrast, our results suggest that talin in biocondensates can induce integrin activation in the absence of ligand. This discrepancy may result from the presence of other key molecules in the cell, such as kindlin, which drastically increases the affinity of talin for the β-integrin tail^60^. Furthermore, previous work has shown that mechanical engagement is not required for integrin activation by talin^64^ and that the dimerized form of talin exhibits significantly enhanced activity over the monomeric state^37,54^ by enhancing cooperation with kindlin and paxillin. This may be the mechanism by which LLPS of talin enhances integrin activation as well. LLPS maintains proteins in a crowded environment, while still allowing them to be dynamic. Thus, it increases their dimerization potential and keeps integrins, talin, and other adaptors enriched in a region, throughout repeated binding and unbinding. This is not dissimilar to other examples of LLPS providing spatial memory through sequestering proteins in a region of a cell such as with P granules.^65,66^ A growing body of evidence^22,23,67^ shows that proteins within focal adhesions are very dynamic, yet this does not cause them to disassemble. While binding of both kindlin and talin to the β-integrin is required for focal adhesion formation, they exist in a dynamic equilibrium where only one is bound in most complexes at any given time.^60^ Similarly, in a biocondensate, at any given time there may be integrin-bound and unbound talins. However, as long as the talins remain at a high concentration in the condensate they can interact with and sequester integrins at the site of the condensate in a matrix-independent manner. This is consistent with our simulations, in which LLPS of talin enables it to compete with other existing adhesions for integrins.

Molecular clutch theory, in which the clutch bonds between the integrins and the ECM are stabilized by traction force, explains the promotion of adhesions around the edge of cells and the general lack of stabilization of our central adhesions in the absence of light activation. Furthermore, our model reveals how biocondensation of talin can occur much more readily at the cell periphery, thereby determining where adhesions form. In the center of our cells, the lack of actin flow and of ventral actin likely prevents the stabilization of central adhesions in the absence of light. Experiments showing that zyxin is recruited to sites of stress fiber strain^68^ and is force sensitive^69^ support this hypothesis. Vinculin, while thought to be an important part of the molecular clutch, paradoxically has been shown not to be essential for focal adhesion growth in response to actin flow^70^. Adhesion growth through increased phase separation may reconcile these observations possibly via talin unfolding or recruitment if FAK or paxillin, leading to increased phase separation. While the central regions of cells where we created IACs generally had little ventral actin, some actin was recruited by the IACs themselves. The molecular pathways of F-actin assembly around central optogenetic IACs are likely triggered by formin activation and/or Arp2/3-based branching^49,71^. However, even though some actin can be nucleated in the middle of the cell, the actin composition there may not contain all the correct forms of actin which have recently been shown to form in layers of distinct composition in IACs^72^.

While the evidence of the role of LLPS in focal adhesions has rapidly increased in recent years, we still have limited evidence regarding the function of LLPS in focal adhesion formation. This work demonstrates that light-induced LLPS can enable *de novo* formation of IACs, maintain integrin activation, and provides a new optogenetic platform to explore adhesion nucleation and function. Our model explains how molecular clutch formation works in combination with LLPS to promote adhesion formation and stabilization in specific areas of the cell. Furthermore, the ability to create new adhesions in a spatiotemporally defined manner is a very powerful technique for studying adhesion dynamics and function. These tools will be invaluable for understanding the roles of specific adhesion states and the role of local adhesion formation. Future studies will be needed to better understand how LLPS is regulated in adhesions throughout their lifetimes and to develop optimized strategies to construct synthetic adhesions based on these principles.

## Supplementary Figures

**Figure S1:**
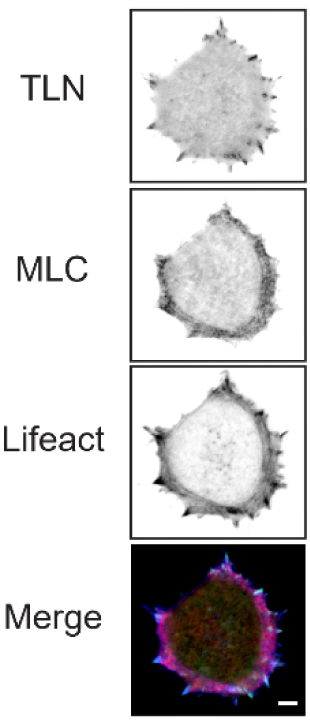
Myosin II and actin distribution in HEK 293T cells. Representative TIRFM images of HEK 293T cells co-expressing GFP-TLN, MLC-mScarlet, and Lifeact-miRFP680 (n=11 cells).

**Figure S2: Related to Figure 2:**
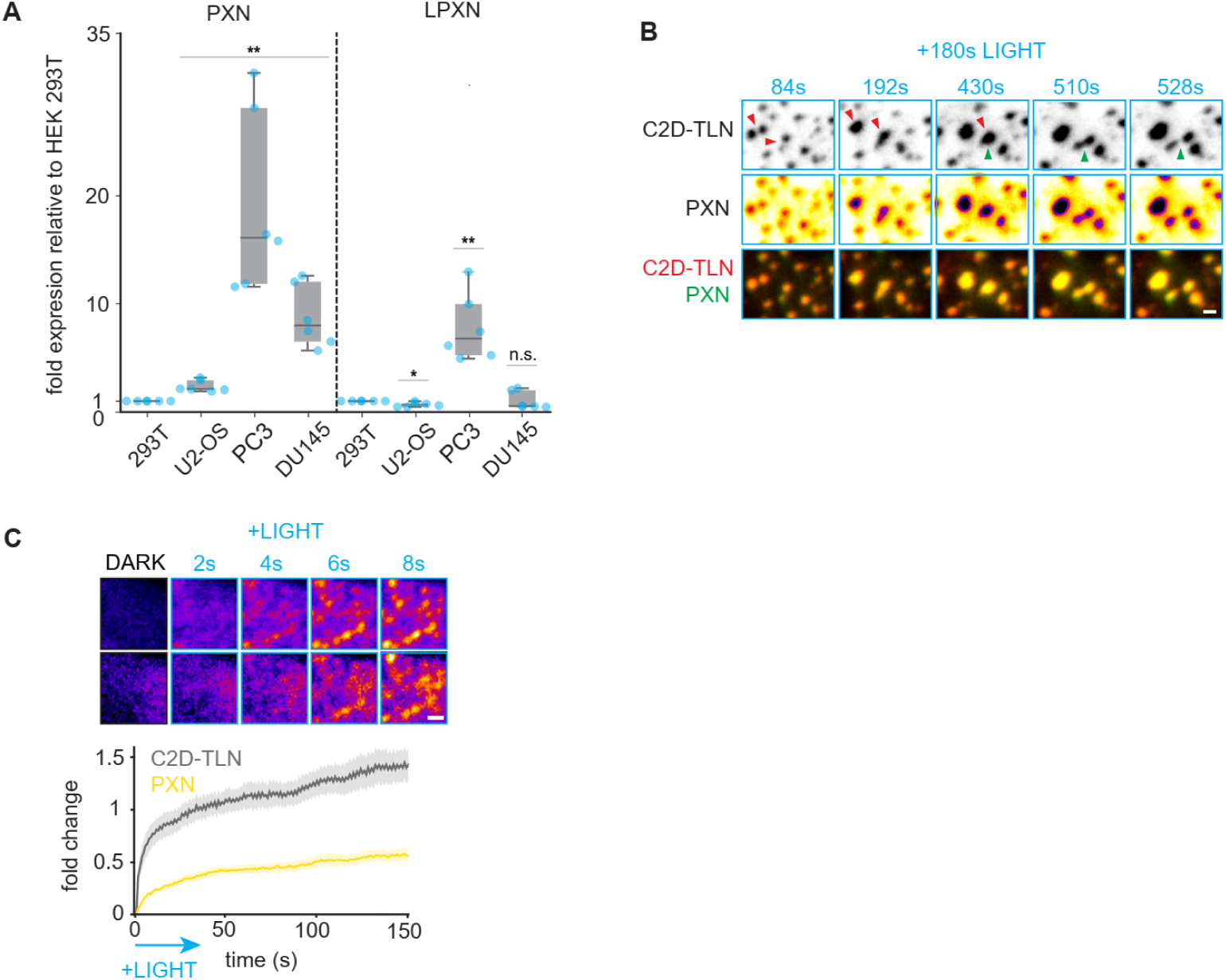
Paxillin enables LLPS of talin and central focal adhesion nucleation. **A)** Expression levels of paxillin and its isoforms quantified by qPCR quantified in several cell lines (N=6, one-sample t-test, μ_0_=1; \**P*<0.05, \*\**P*<0.01). **B)** Montage of droplet fusions and fissions of FusionRed-C2D-TLN trigged by global light activation in HEK 293T cells with the additional expression of PXN-miRFP680 on the polylysine-coated glass. Red arrows indicate droplets undergoing fusion and green arrows indicate fission. See also Movie S7. **C) (**Top) montage of the first few seconds of PXN-miRFP680 and FusionRed-C2D-TLN accumulation after light illumination in HEK 293T cells seeded on polylysine and (bottom) quantification (n=7 cells, error bars ± S.E.M.). Scale bars: 1μm.

**Figure S3: Related to Figure 3:**
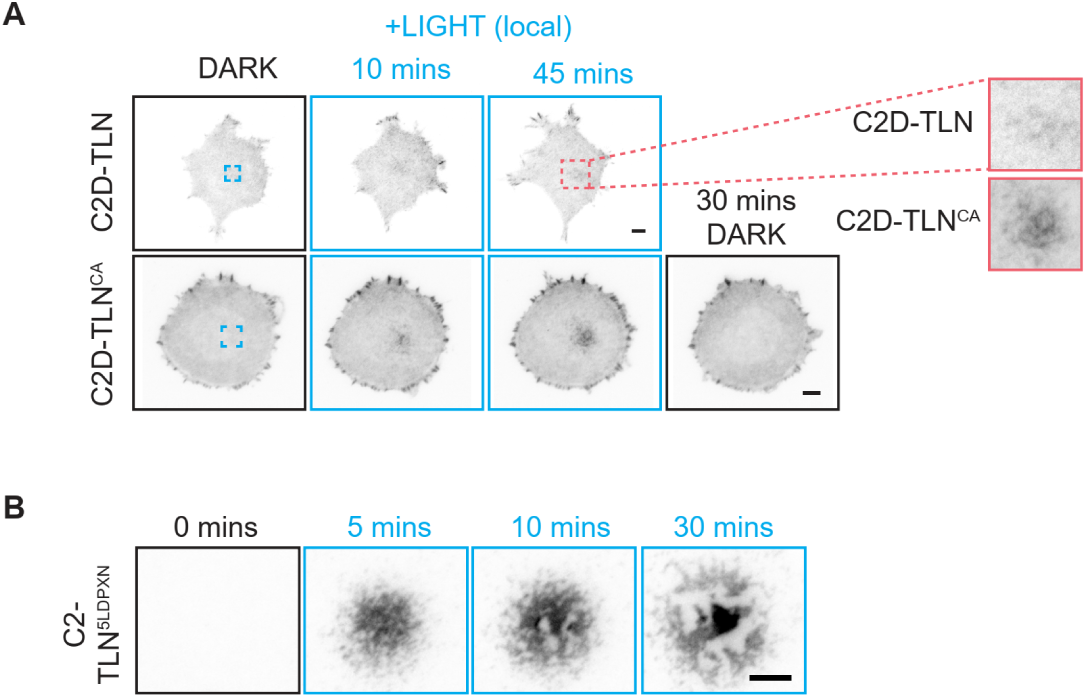
LLPS of talin allows for adhesion nucleation. **A)** Montage of cells expressing FusionRed-C2D-TLN or FusionRed-C2D-TLN^CA^ exposed to central illumination in the blue box for 45 minutes before being allowed to recover for another 30 mins. Zooms of central regions after 45 minutes activation shown to the right. **B)** Montage showing the spatiotemporal separation of distinct phases of FusionRed-C2D-TLN^5LDPXN^ during central illumination. Scale bars: 5μm

**Figure S4: Related to Figure 4:**
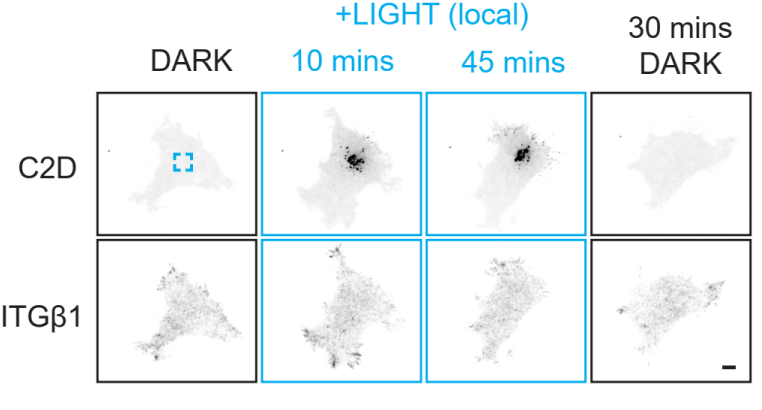
Phase separation of talin partitions and activates integrins. Montage of a cell co-expressing ITGβ1-miRFP680 and FusionRed-C2D a, exposed to central illumination in the blue box for 45 minutes before being allowed to recover for another 30 mins. Scale bar: 5μm

**Figure S5: Related to Figure 6:**
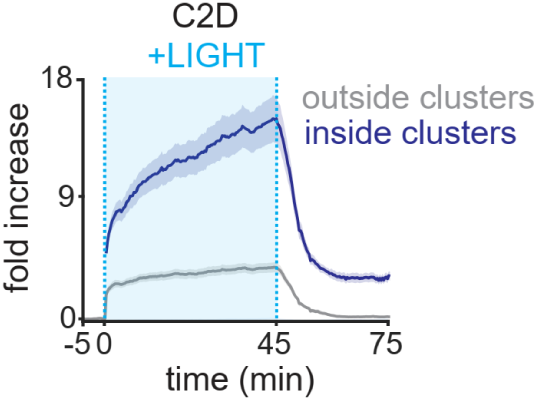
Model of adhesion nucleation by LLPS. **A)** Intensity on the membrane of C2D inside and outside C2D droplets upon light activation used to estimate kBT for C2D.

## Supplementary Movies

**Movies can be found at** https://www.johnsonlaboratory.org/supplemental-data

<u>Movie S1: Global illumination of C2-TLN, related to Figure 1B</u>

<u>Movie S2: Global illumination of C2D-TLN, related to Figure 1B</u>

<u>Movie S3: Local illumination of C2D-TLN, related to Figure 1E</u>

<u>Movie S4: Global illumination of C2D, related to Figure 2A</u>

<u>Movie S5: Global illumination of C2D-TLN with PXN on FN ,related to Figure 2A</u>

<u>Movie S6: Global illumination of C2D-TLN with PXN on PL, related to Figure 2A</u>

<u>Movie S7: Droplet fusion/fission during global illumination of C2D-TLN with PXN on PL, related to Figure S2B</u>

<u>Movie S8: Central activation of modified talins, related to Figure 3B</u>

<u>Movie S9: Central activation of C2D-TLN with VCL or VCL^DR^, related to Figure 3E</u>

<u>Movie S10: Central activation of C2-TLN^5LDPXN^ with ITGβ1, related to Figure 4C</u>

<u>Movie S11: Central activation of C2D-TLN^5LDPXN^ with Lifeact, related to Figure 5A</u>

<u>Movie S12: Central and edge activation of C2D-TLN^3CA^ with ZYX, related to Figure 5C</u>

<u>Movie S13: Model of molecular clutches for C2D-TLN central activation, related to Figure 6D-F</u>

<u>Movie S14: Model of molecular clutches for C2D-TLN^3CA^ central activation, related to Figure 6D-F</u>

## Author Contributions

H.E.J. conceived the project M.X., C.Y., and H.E.J. designed experiments, M.X. performed experiments. M.X., M.C., J.C. and T.C.L cloned constructs. T.L.L performed preliminary experiments, A.K.E. created the model and ran the computations. H.E.J analyzed data. H.E.J and A.K.E created illustrations, H.E.J. wrote the manuscript with input from all authors.

## Acknowledgement

We would like to thank Dr. Chenyu Mao (Tsinghua Shenzhen International Graduate School) for sharing pcDNA3.1 backbone plasmids, Haozhe Zhang (SUSTech) for sharing PC-3 and DU145 cell lines. This work was supported by the Research Grant Council of Hong Kong, CRF C7070-22EF (C.H.Y.), HKU Seed Grant for basic research #2202100793 (H.E.J.), as well as start-up funds from HKU (H.E.J.) and SZBL (A.K.E.).

